# Imputation and polygenic score performances of human genotyping arrays in diverse populations

**DOI:** 10.1101/2022.06.14.496059

**Authors:** Dat Thanh Nguyen, Trang Tran, Mai Tran, Khai Tran, Duy Pham, Nguyen Thuy Duong, Quan Nguyen, Nam S. Vo

**Affiliations:** Center for Biomedical Informatics, Vingroup Big Data Institute, Hanoi, Vietnam; Institute for Molecular Bioscience, University of Queensland, Brisbane, Australia; Institute of Genome Research, Vietnam Academy of Science and Technology, Hanoi, Vietnam; College of Engineering and Computer Science, VinUniversity, Hanoi, Vietnam

## Abstract

Regardless of the overwhelming use of next-generation sequencing technologies, microarray-based genotyping combined with the imputation of untyped variants remains a cost-effective means to interrogate genetic variations across the human genome. This technology is widely used in genome-wide association studies (GWAS) at bio-bank scales, and more recently, in polygenic score (PGS) analysis to predict and to stratify disease risk. Over the last decade, human genotyping arrays have undergone a tremendous growth in both number, and content making a comprehensive evaluation of their performances became more important. Here, we performed a comprehensive performance assessment for 23 available human genotyping arrays in 6 ancestry groups using diverse public, and in-house datasets. The analyses focus on performance estimation of derived imputation (in terms of accuracy and coverage) and PGS (in term of concordance to PGS estimated from whole genome sequencing data) in three different traits and diseases. We found that the arrays with a higher number of SNPs are not necessarily the ones with higher imputation performance, but the arrays that are well-optimized for the targeted population could provide very good imputation performance. In addition, PGS estimated by imputed SNP array data is highly correlated to PGS estimated by whole genome sequencing data in most of cases. When optimal arrays are used, the correlations of key PGS metrics between two types of data can be higher than 0.97, but interestingly, arrays with high density can result in lower PGS performance. Our results suggest the importance of properly selecting a suitable genotyping array for PGS applications. Finally, we developed a web tool that provide interactive analyses of tag SNP contents and imputation performance based on population and genomic regions of interest. This study would act as a practical guide for researchers to design their genotyping arrays-based studies. The tool is available at: https://genome.vinbigdata.org/tools/saa/

## 1 INTRODUCTION

Over the last decade, low-cost, robust genotyping platforms and large-scale genome variation projects such as the 1000 Genomes Project (Auton et al., 2015) have facilitated genome-wide association studies (GWAS) on numerous human phenotypes, ranging from height to diseases (Bycroft et al., 2018). To date, thousands of DNA loci that are significantly associated with complex traits and diseases have been discovered (Buniello et al., 2019). Among numerous possible applications of GWAS results, disease risk prediction is rapidly gaining broad interest recently (Torkamani et al., 2018; Lewis and Vassos, 2020; Lambert et al., 2019). A polygenic score (PGS) or polygenic risk score (PRS) is an estimate of an individual’s genetic liability to a trait or disease, calculated based on their genotype profile and relevant GWAS data (Choi et al., 2020). In its most common form, a PGS is computed as the sum of allele count of risk alleles (0, 1, or 2) that are weighted by its effect size (i.e. log odd ratio or beta coefficient) of hundreds-to-thousands of associated SNPs. The outcome is a single score that aggregate each individual’s genetic loading proportional to the risk of a given disease or a quantitative trait (Lambert et al., 2019). Although clinical utility of PGS has yet to be established, recent works have suggested that PGS may be used for disease risk stratification that potentially facilitates early disease detection, assists in diagnosis or informs treatment choices (Torkamani et al., 2018; Lewis and Vassos, 2020). For example, PGS of coronary artery disease, type 2 diabetes, and breast cancer at the top 8, 3.5, and 1.5% are risk equivalent to a monogenic mutation risk that confers an odd ratio of 3 (Khera et al., 2018).

Similar to GWAS analysis, PGS can be derived from various types of genotyping data such as those obtained by single-nucleotide polymorphism (SNP) microarrays or whole genome sequencing (WGS). While WGS is attractive by the ability to interrogate variations across the entire human genome, SNP arrays are the dominant assays to obtain genetic data for PGS calculation. They come up with several advantages such as cost-effectiveness and light computational requirement which are preferable for population-sale screening, where PGS would be most useful (Chen et al., 2020). Because the coverage of SNP arrays is typically limited to lower than a million SNPs, a procedure involving haplotye phasing and genotype imputing of missing sites is usually employed to add more genotyping information that can increase powers of these genetic studies (Howie et al., 2012; Marchini and Howie, 2010; Choi et al., 2020). The imputation performance is affected by three main factors, including algorithms of choice (Das et al., 2016), imputation reference panels (Huang et al., 2015; McCarthy et al., 2016), and the SNP array designs (Nelson et al., 2013).

In principle, genotyping SNP arrays are designed by selecting a set of SNPs, commonly reffered to as “tag SNPs”, which maximize coverage of ungenotyped DNA variants through associations between these alleles in the population (known as linkage disequilibrium, LD) (Gibbs et al., 2003; Carlson et al., 2004). Based on the target population, human genotyping SNP arrays can be classified into three categories that optimized for global, super population or specific to targeted populations. In the early phase of development, genotyping SNP arrays were focused on common genetic variations of the whole world population (minor allele frequency, MAF, of 0.10 or greater) based on the HapMap catalog (Consortium et al., 2007). The second generation of SNP arrays were designed to cover variants with MAF as low as 0.01 by providing SNP arrays specifically for European, East Asian, African American, and Latino race/ethnicity populations based on the 1000 Genomes Project (1KGP) catalog (Hoffmann et al., 2011a,b). However, the fact that the majority of human genetic variants are rare and population-specific demands customizing SNP arrays to improve over those designed for global or super populations (Consortium et al., 2015; Tam et al., 2019). Indeed, population-specific genotyping arrays such as the UK Biobank Axiom Array (Bycroft et al., 2018), the Axiom-NL Array (Ehli et al., 2017), the Japonica and Japonica NEO Arrays (Kawai et al., 2015; Sakurai-Yageta et al., 2020), and the Axiom KoreanChip (Moon et al., 2019) have been developed on top of the many existing commercial arrays. These arrays are not only optimized for genomic coverage based on their unique variant catalogs, but also include a large amounts of functional variants. For example, the Axiom KoreanChip contains more than 200,000 nonsynonymous loci and the new Japonica NEO Arrays is designed with abundant disease risk variants (Moon et al., 2019; Sakurai-Yageta et al., 2020).

The development of customized arrays together with commercial arrays provided by genotyping platform producers result in a large number of genotyping arrays. Each of these arrays has specific properties and contents, and thus, there is an urgent demand for a systematic guideline to determine which array best suits specific research questions and populations. Although there are SNP array comparative studies, they are either not updated with the many recent arrays (Nelson et al., 2013; Ha et al., 2014), or limited in only testing for a small set of populations, and some studies focused on LD coverage (Ha et al., 2014; Verlouw et al., 2021) that may not be relevant to current imputation practice for use in association studies and PGS analysis (Marchini and Howie, 2010; Choi et al., 2020). Moreover, although PGS is gaining increasing attention, practical evaluation of performance for PGS applications by current genotyping arrays is still lacking. Here, we provide a comprehensive evaluation of imputation-based genomic coverage (Lindquist et al., 2013; Nelson et al., 2013) and PGS performance of 23 human genotyping arrays in diverse populations. These analyses are intended to be a practical guide for researchers in selecting the most suitable genotyping array for their genetic studies.

## 2 MATERIALS AND METHODS

### 2.1 Genotyping arrays

In this study, we benchmarked 23 different human genotyping arrays including 14 arrays from Illumina and 9 arrays from Affymetrix. The examined arrays contain the numbers of tag SNPs (array size) ranging from approximately 300,000 (Infinium HumanCytoSNP-12 v2.1) up to more than 4,300,000 (Infinium Omni5 v1.2). They can be classified as old arrays such as the Genome-Wide Human SNP Array 6.0; population specific optimized arrays such as Axiom UK Biobank Array and Axiom Japonica Array NEO; multiple population optimized arrays such as Infinium Multi-Ethnic Global v1.0 and Infinium Global Diversity Array v1.0; cytogenetics and cancer applications optimized arrays such as Infinium CytoSNP-850K v1.2. Recently developed arrays include Infinium Global Screening Array v3.0, Axiom Precision Medicine Research Array, and Axiom Precision Medicine Diversity Array. Manifests of the 23 examined arrays were obtained from respective manufacturers’ websites. Genomic positions were further harmonized to the UCSC hg38 reference genome coordinate with CrossMap v0.2.6 for those require lifted over (Zhao et al., 2014). Details and component statistics of these arrays are shown in Table 1.

**Table 1.**
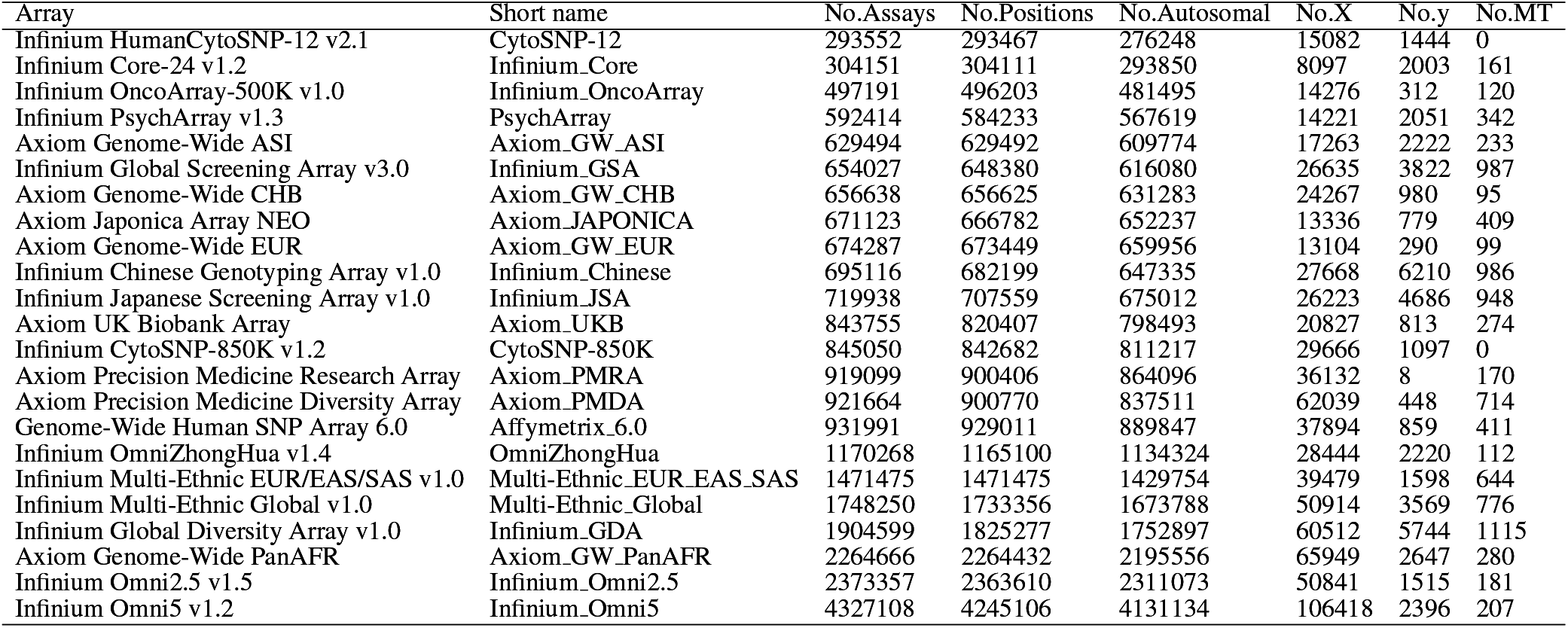
Details of 23 human genotyping arrays used in this study.

### 2.2 Genomic datasets and pipelines

An overview of our evaluation pipeline is presented in Figure 1. In brief, the phased genomic data in Variant Call Format (VCF) of 2,504 and 1,008 unrelated individuals from the 1000 Genomes Project samples that were re-sequenced by New York Genome Center (1KGP) (Byrska-Bishop et al., 2021) and the 1000 Vietnamese Genomes Project (1KVG) (Tran et al., 2022) were used to estimate imputation-based coverage and PGS performance of 23 different genotyping arrays by the 10-fold cross-validation approach. Specifically, genomic datasets were randomly divided into 10 batches (equally distributed across populations in the 1KGP dataset). In each turn, one batch was used as the test set and the remaining samples as the reference set. For each array, variants in the test set with the same position as variants on the array were extracted with vcftools v0.1.17 (Danecek et al., 2011) and phasing information was removed to generate the pseudo SNP array genotyped data, while variants in reference data were used as the pre-phasing and imputation reference panel. The pre-phasing and imputation were performed with SHAPEIT v4.1.3 (Delaneau et al., 2019) and Minimac4 v1.0.2 (Das et al., 2016) respectively. Finally, the imputed genotyping data of 10 batches were combined to estimate imputation and PGS performance according to their populations, including 504, 503, 489, 661, 347, and 1,008 individuals in East Asian (EAS), European (EUR), South Asian (SAS), African (AFR), American (AMR) and Vietnamese (VNP) populations, respectively. This approach is similar to the strategy used previously to estimate imputation-based genomic coverage (Lindquist et al., 2013; Nelson et al., 2013; Nguyen et al., 2021).

**Figure 1.**
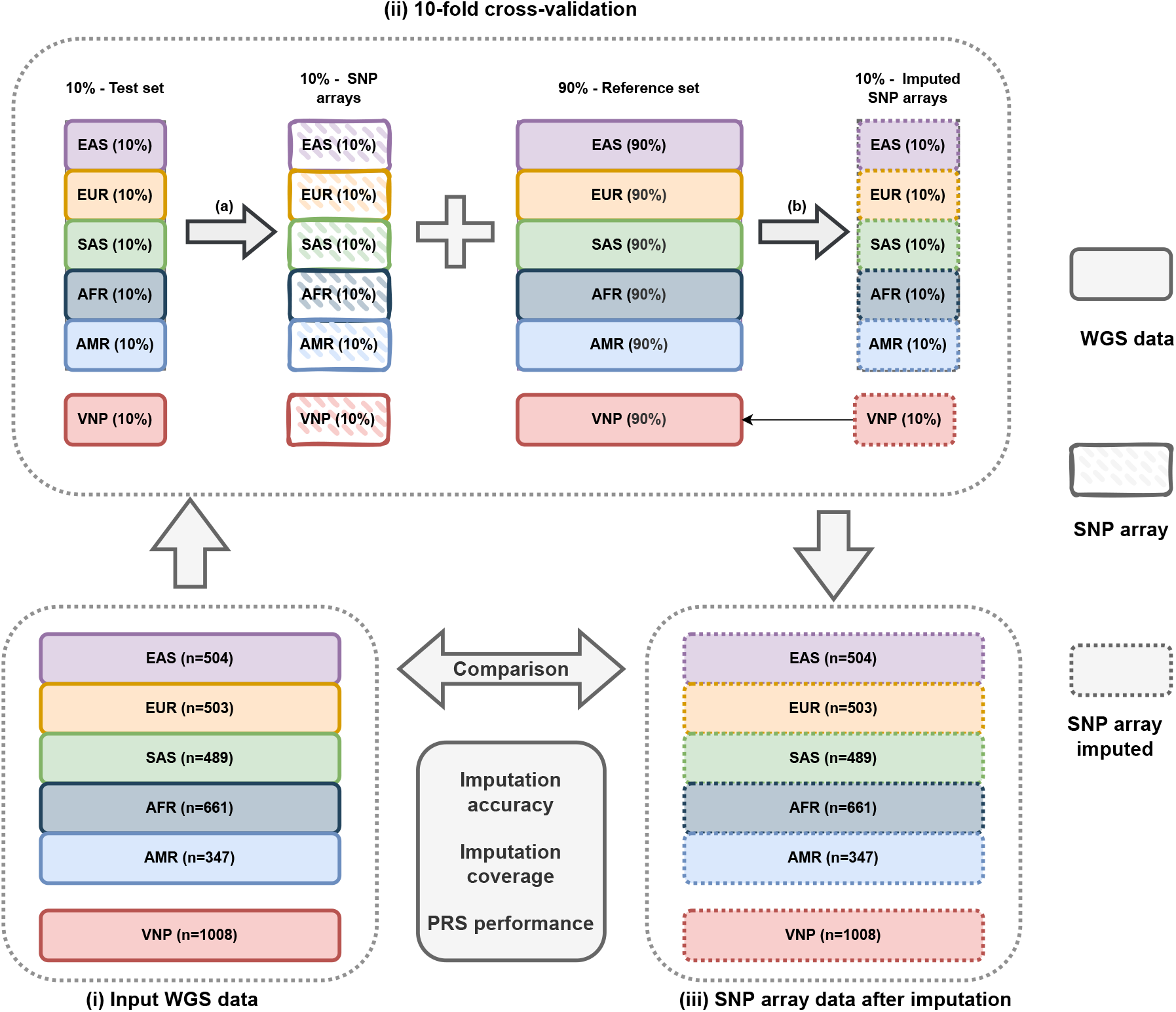
Overview of evaluation pipeline. (i) Two input genetic datasets, including the 1KGP and 1KVG were randomly divided into 10 batches that are equally distributed by populations. (ii) 10-fold crossvalidation procedure. In each turn, variants of 10% samples were extracted based on arrays’ manifest to generate simulated array genotyping data (arrow a) as input for phasing and imputation with the remaining 90% samples used as the reference set to generate the imputed SNP array data (arrow b). (iii) SNP array data after imputation. Imputed SNP array data of 10 batches were merged according to populations after 10-fold cross-validation, and were then benchmarked by treating the input WGS data as the golden standard.

### 2.3 Imputation performance evaluation

Both GWAS and PGS often require genotype imputation that involves prediction of untyped variants in the genome. While GWAS benefits from the boosting the number of imputed SNPs that can be tested for association (Marchini and Howie, 2010), computation of PGS is conducted by summing the product of risk allele count (0, 1, or 2) and its effect size derived from the GWAS. Thus, imputation performance is expected to play a key role in PGS derivation. Here, we focus on imputation *r*^2^ metric although there are several other criteria that can be used to assess imputation performance such as allele concordance (Nelson et al., 2013), imputation quality (Verlouw et al., 2021), LD coverage (Barrett and Cardon, 2006). We choose imputation *r*^2^ as the evaluation metric for following reasons. First, it is more relevant to the context of GWAS and PGS analysis because the imputation *r*^2^ at a given variant is proportional to its *χ*^2^ statistic that results from an association test (Pritchard and Przeworski, 2001; Chapman et al., 2003; Marchini, 2019; Li et al., 2021). This leads to the interpretation that an increase in mean imputation *r*^2^ at genome wide scale directly corresponds to the increase of statistical power (Pritchard and Przeworski, 2001; Li et al., 2021). Second, it is less sensitive to allele frequency than concordance (Nelson et al., 2013). Third, it incorporates imputation uncertainty by using expected allele dosage rather than the most likely genotype (Nelson et al., 2013). Finally, imputation *r*^2^ can be computed on a site-by-site basis, which enables more detailed evaluation than at the allele frequency level (Li et al., 2021). In this evaluation setting, we treated genotypes derived from WGS datasets as gold-standard. Imputation performance is measured as imputation *r*^2^ that is SNP-wise squared Pearson’s correlation between the imputed dosages and the WGS genotypes, and imputation coverage that is defined as proportion of SNPs with imputation *r*^2^ passing the cut-off of 0.8. These metrics were stratified into three minor allele frequency (MAF) bins, including (0-0.01], (0.01-0.05], (0.05-0.5]. To reduce the data noise, variants with allele count ¡ 2 are excluded in the bin of (0-0.01]. Of note, the MAF bin of (0.01-0.5], which is the most common cutoff for GWAS and PGS analysis, was also considered in the analysis (Marees et al., 2018; Choi et al., 2020).

### 2.4 PGS performance assessment

Instead of using pre-tuned PGS models as other studies (Li et al., 2021; Chen et al., 2020)), in this study PGS was computed with a standard P+T (Prunning and Thresholding) approach implemented in PRSice-2 (Choi and O’Reilly, 2019). The main reason for using this approach is that we tried to mimic the real-life practice of PGS analysis that involves running a PGS computational method with multiple parameters and selecting the best one (Choi et al., 2020). Another reason was that using pre-built PGS models may introduce a potential bias for some specific arrays as they were used in tuning, and we tried to avoid training using the same array twice. Using summary statistics for three phenotypes, namely height, body mass index (BMI), and type 2 diabetes (T2D), obtained from previous GWAS meta analyses (Yengo et al., 2018; Xue et al., 2018), a PGS for an individual *i* was calculated as:

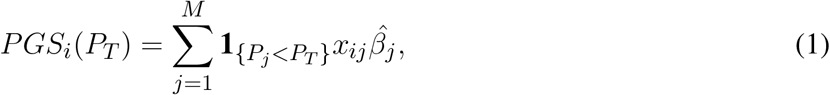

where *P*_*T*_ is the p-value threshold values (5e-08, 1e-07, 1e-06, 1e-05, 0.0001, 0.001, 0.01, 0.1, 0.2, 0.3, 0.5, and 1); *M* is number of SNPs after clumping with *“–clump-kb 250kb”* and *“–clump-r2 0*.*1”*; *x*_*ij*_ and 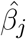 is the allele count and the marginal effect size derived from GWAS summary statistics of *SNP*_*j*_.

Similar to imputation performance evaluation, we treat PGSs derived from WGS as the “gold-standard”. PGSs derived from 23 different SNP arrays were evaluated using Pearson’s correlation and absolute percentile differences compared to the gold-standard under the same PRSice-2 parameter settings. In addition, absolute difference of PGS percentile ranking generated by array-imputed and the gold-standard was also evaluated.

## 3 RESULTS

### 3.1 Imputation performance

Overall, we found two main factors affecting the imputation accuracy and imputation coverage, including array sizes and population-specific optimization. The Infinium Omni2.5 v1.5 and Infinium Omni5 v1.2 with approximately 2.4 and 4.3 minion tag SNPs yielded the highest imputation performance. In contrast, low density SNP arrays with approximately 300,000 tag SNPs such as Infinium HumanCytoSNP-12 v2.1 and Infinium Core-24 v1.2 obtain the poorest imputation performance in all all six examined populations. At the MAF bin of (0.01-0.5], the Infinium Omni5 v1.2 yielded the mean imputation accuracy *r*^2^ of 0.9032, 0.9144, 0.8644, 0.9176, 0.8873, 0.9499 and the imputation coverage of 0.8721, 0.8813, 0.8019, 0.8885, 0.8344, 0.9207 while the Infinium HumanCytoSNP-12 v2.1 obtained 0.6682, 0.7708 0.7112, 0.7608 0.7218, 0.8635 for mean imputation accuracy *r*^2^ and 0.4031, 0.6265, 0.5879, 0.6297, 0.5731, 0.7655 for imputation coverage in six populations AFR, AMR, EAS, EUR, SAS, and VNP respectively. Details are reported in Figure 2 and Table 2, 3.

**Table 2.**
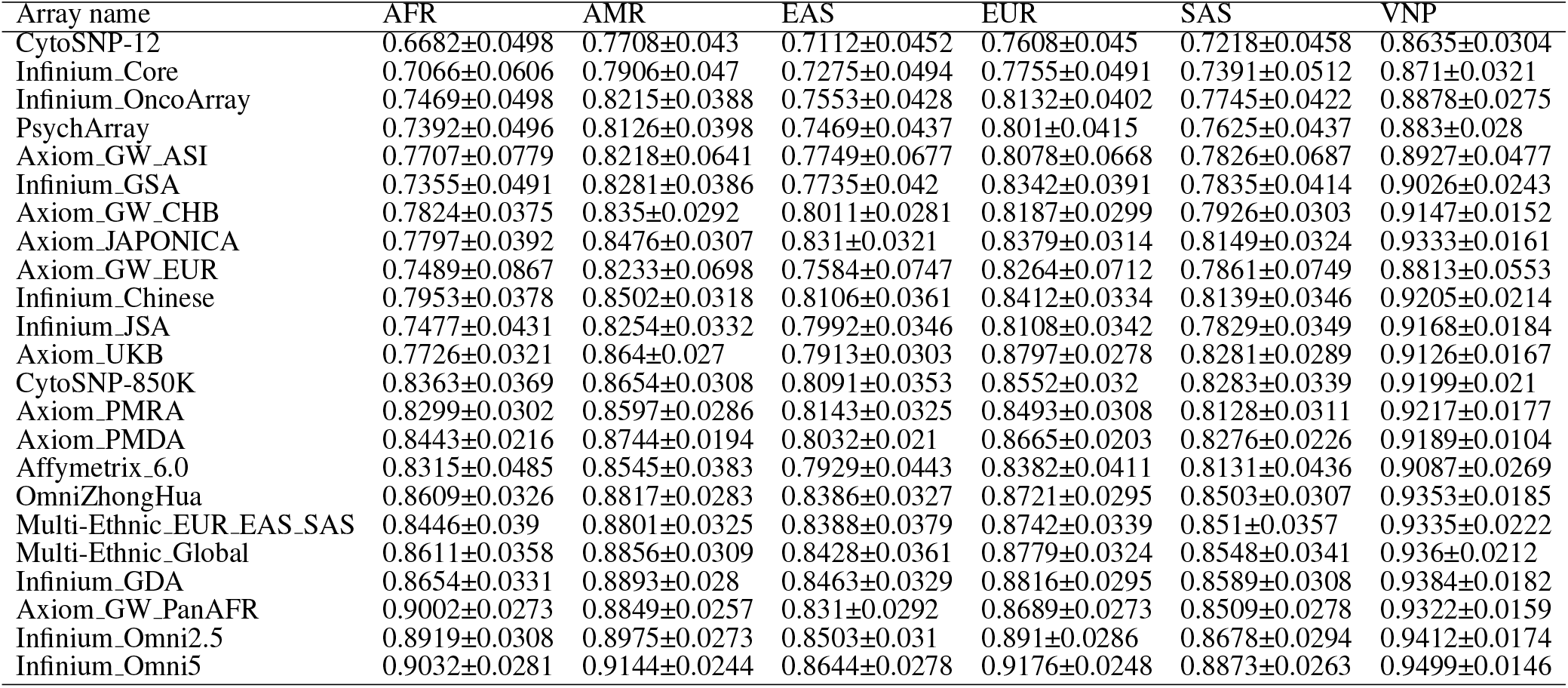
Mean and the standard deviation of imputation accuracy *r*^2^ measured in 22 autosomes at the MAF bin of (0.01-0.5].

**Table 3.**
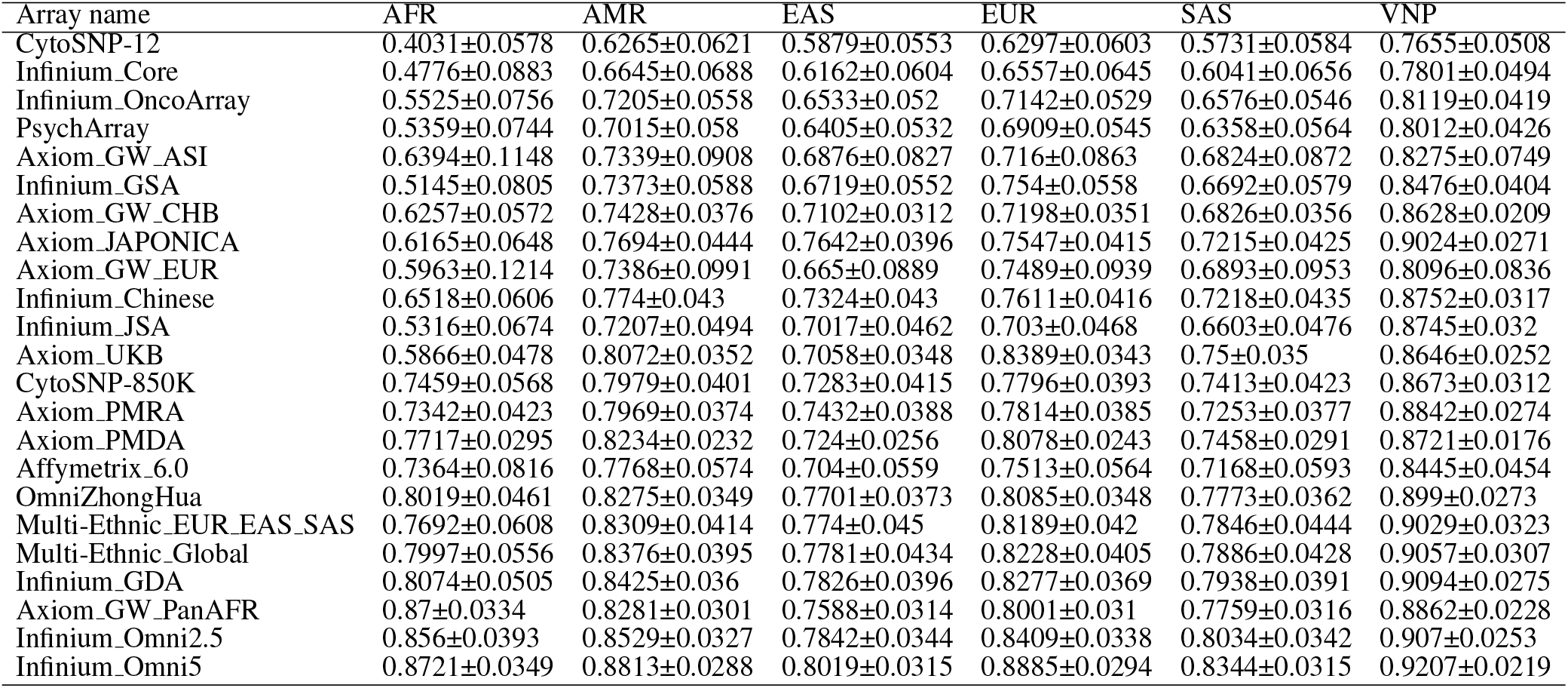
Mean and standard deviation of imputation coverage (defined by the proportion of variants with *r*^2^*≥* 0.8 over total number of variants in each chromosome) measured in 22 autosomes at the MAF bin of (0.01-0.5].

**Figure 2.**
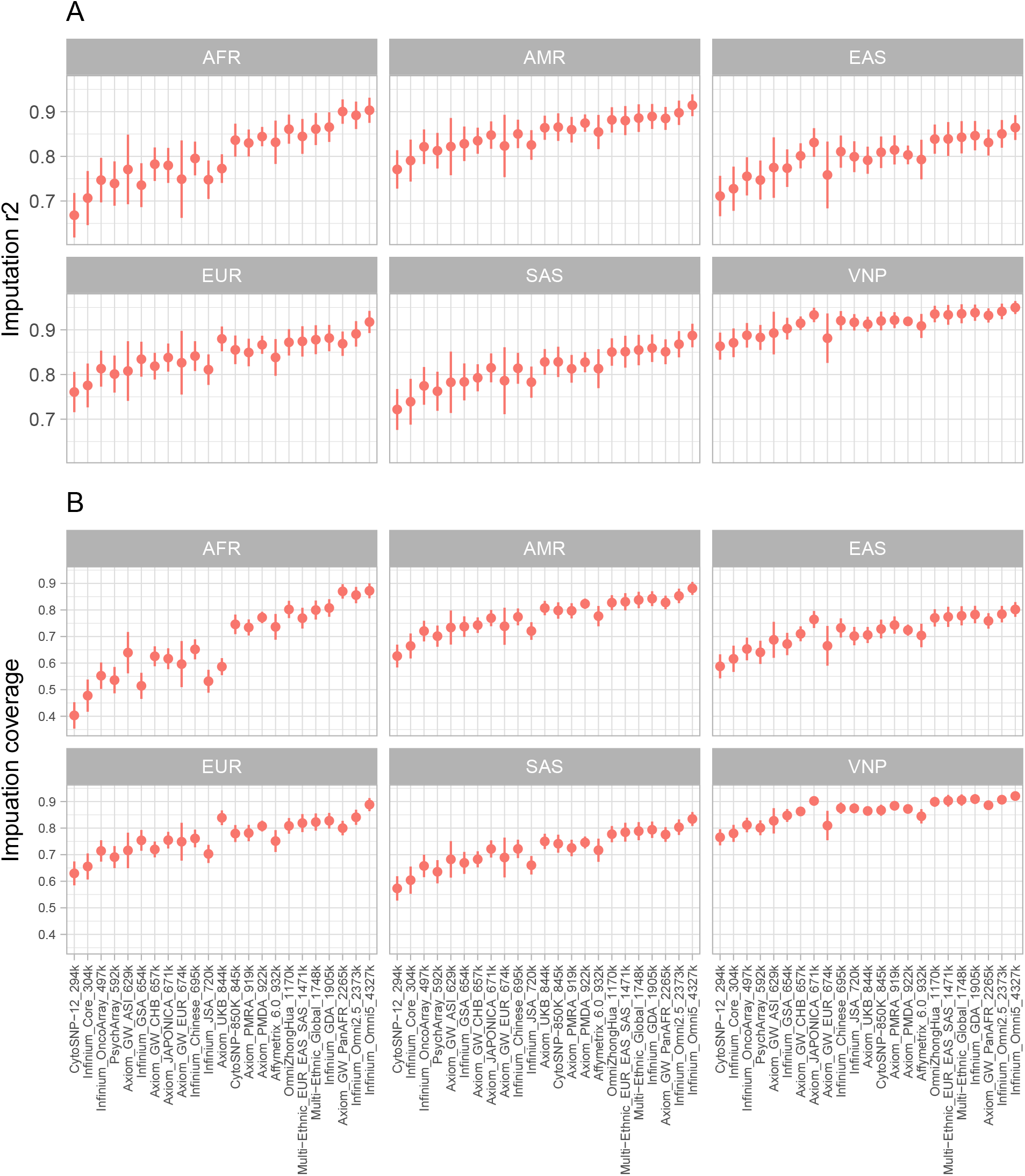
A. Mean imputation *r*^2^, and B. Imputation coverage across 22 autosomes of 23 SNP arrays in the MAF bin of (0.01-0.5]. The dots and the vertical lines present the mean and the standard deviation of imputation accuracy, and imputation coverage values in 22 autosomes respectively.

Regarding population optimization, imputation performance are generally better for those arrays optimized specifically for their closely related populations. The Axiom UKB, which is optimized for the British population, performs superior for the EUR than most other arrays, except for the ultra high density arrays Infinium Omni2.5 v1.5 and Infinium Omni5 v1.2. In detail, at the MAF bin of (0.01-0.5], The Axiom UKB with size of 844k SNPs obtained the mean imputation coverage of 0.8389 which was higher than globally optimized, higher density arrays such as Axiom PMRA (919k), Axiom PMDA (922k), Affymetrix 6.0 (932k), Multi-Ethnic Global (1784k), and Infinium GDA (1905k), with imputation coverage of 0.7814, 0.8078, 0.7513, 0.8228, 0.8277, respectively and lower 0.8409, and 0.8885 that were obtained by Infinium Omni2.5 v1.5 and Infinium Omni5 v1.2 arrays with 2373k and 4327k SNPs. Similarly, the Axiom JAPONICA (671k) which was designed for Japanese population also performed well against global optimized, higher-density arrays. These two arrays yielded mean imputation accuracy of 0.831, 0.9333 and imputation coverage of 0.7642, and 0.9024 in EAS and VNP populations. These performances were higher than those of multi-ethics SNP arrays, even with higher density including Axiom PMRA (919k), Axiom PMDA (922k), Affymetrix 6.0 (932k) as showed in Figure 2 and Table 2, 3. Notably, the Chinese GWAS array, OmniZhongHua, was also out performed in the EAS and VNP populations. Regarding the AFR population, an optimized population that is Axiom GW PanAFR with 2265k SNPs. The performance of this array is nearly equivalent the Infinium Omni5 v1.2 array with 4327k SNPs (0.9002 versus 0.9032 for mean imputation accuracy, and 0.8700 versus 0.8721 interns of imputation coverage). There were no SNP arrays with superior performances in the two remaining populations (AMR and SAS), although the Axiom UKB and the Axiom GW AS obtained slightly better performance than other arrays with the same size, when applied for the AMR and SAS populations. In this case, we focused on the MAF bin of (0.01-0.5] as this is most common cutoff allele frequency in both GWAS and PGS analysis (Visscher et al., 2017; Choi et al., 2020). However, the results are also generalized for other bins as shown in Figure S.1 and Table S.1-6.

### 3.2 PGS performance

We evaluated PGS performance of these arrays based on two criteria: (i) Pearson’s correlation of PGSs estimated by using imputed SNP array data compared to the PGSs estimated by using WGS data - hereafter we refer as PGS correlation for short, (ii) absolute difference of percentile ranking (ADPR) between PGSs generated by array-imputed and gold standard WGS. Both comparisons are set under various p-value cutoffs that allows us unbiased evaluate PGS performance of these arrays. In general, we found that PGS performance were highly concordant with imputation performance, i.e. SNP arrays with better imputation performance showed higher correlation with WGS PGSs and less ADPR than the arrays with poor imputation performance.

The summary results of Pearson’s correlation values of PGSs from 23 genotyping SNP arrays for three different phenotypes are shown in Figure 3 and in Tables S.7-9. In general, all examined arrays yielded high PGS correlations. Notably, the vast of majority PGS correlations ranged from 0.90 to 0.99, except for the Infinium HumanCytoSNP-12 v2.1 which had the lowest values. Interestingly, when optimal arrays for populations were used, the PGS correlation to WGS was higher than 0.95. The PGS correlation patterns were also highly concordant in all three evaluated traits with comparable performances. As expected, SNP arrays with larger sizes had higher PGS correlations. The lowest performer was the Infinium HumanCytoSNP-12 v2.1 with the correlation of 0.8731 in the height phenotype in the AFR population while the highest performance was obtained by the Infinium Omni5 v1.2 with the correlation higher than 0.99 in all populations and traits. We also examined the deviation of PGS correlation in various p-value settings. The results showed that SNP array with lower PGS correlation had higher PGS correlation standard deviation than the high performance arrays, A possible explanation for this observation is the PGSs estimated from low imputation performance are more vulnerable by the random pruning process than the high imputation performance arrays (Choi and O’Reilly, 2019). Notably, we also observed the higher standard deviations of PGS correlation in EAS than other populations.

**Figure 3.**
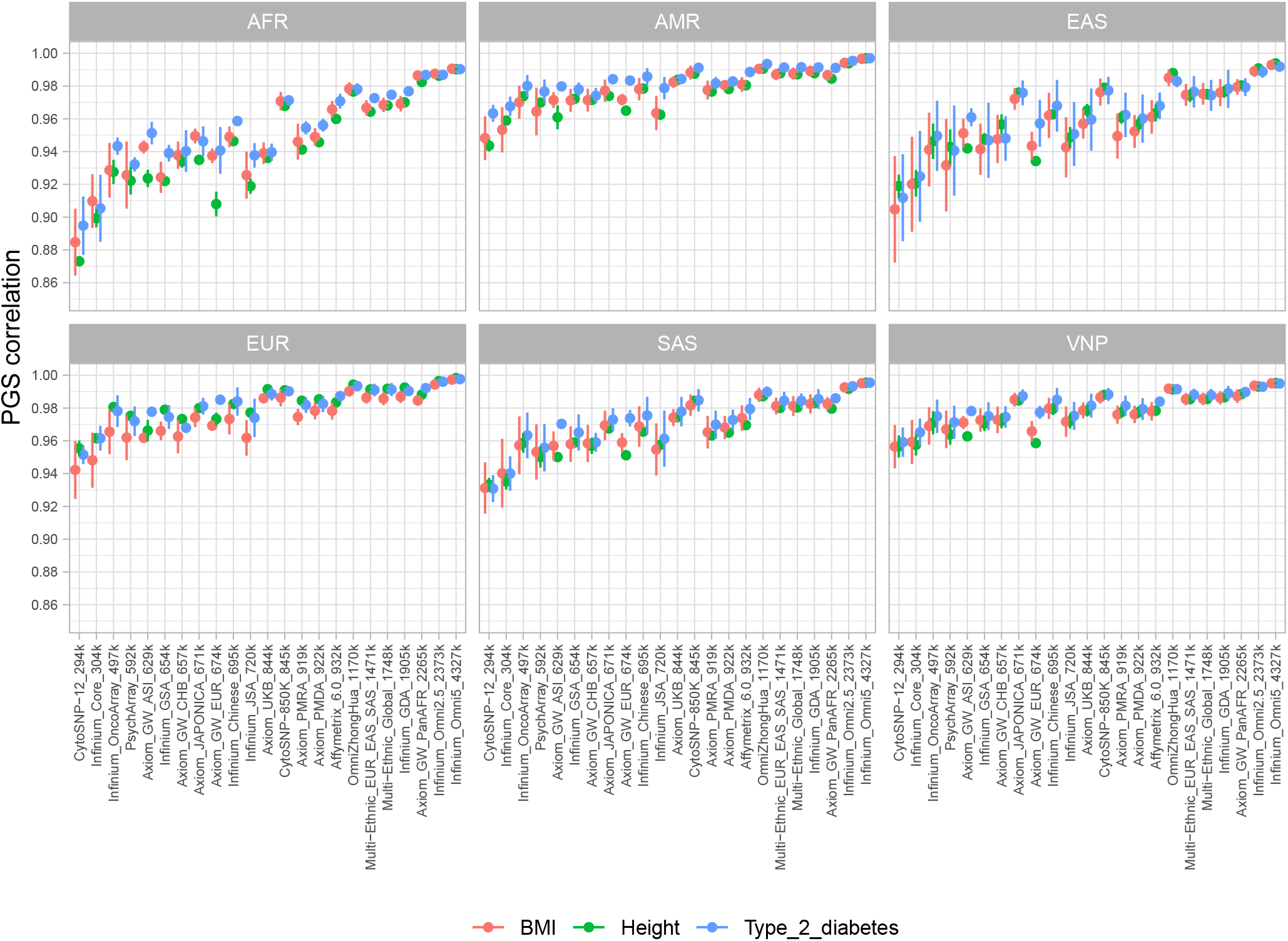
Correlations between PGSs estimated from imputed genotyping data of 23 SNP arrays and PGSs estimated from WGS in six different populations with three phenotypes including height, BMI, and type 2 diabetes. The dots and the vertical lines present the mean and standard deviation of PGS correlation at various p-value settings including 5e-08, 1e-07, 1e-06, 1e-05, 0.0001, 0.001, 0.01, 0.1, 0.2, 0.3, 0.5, and 1.

**Figure 4.**
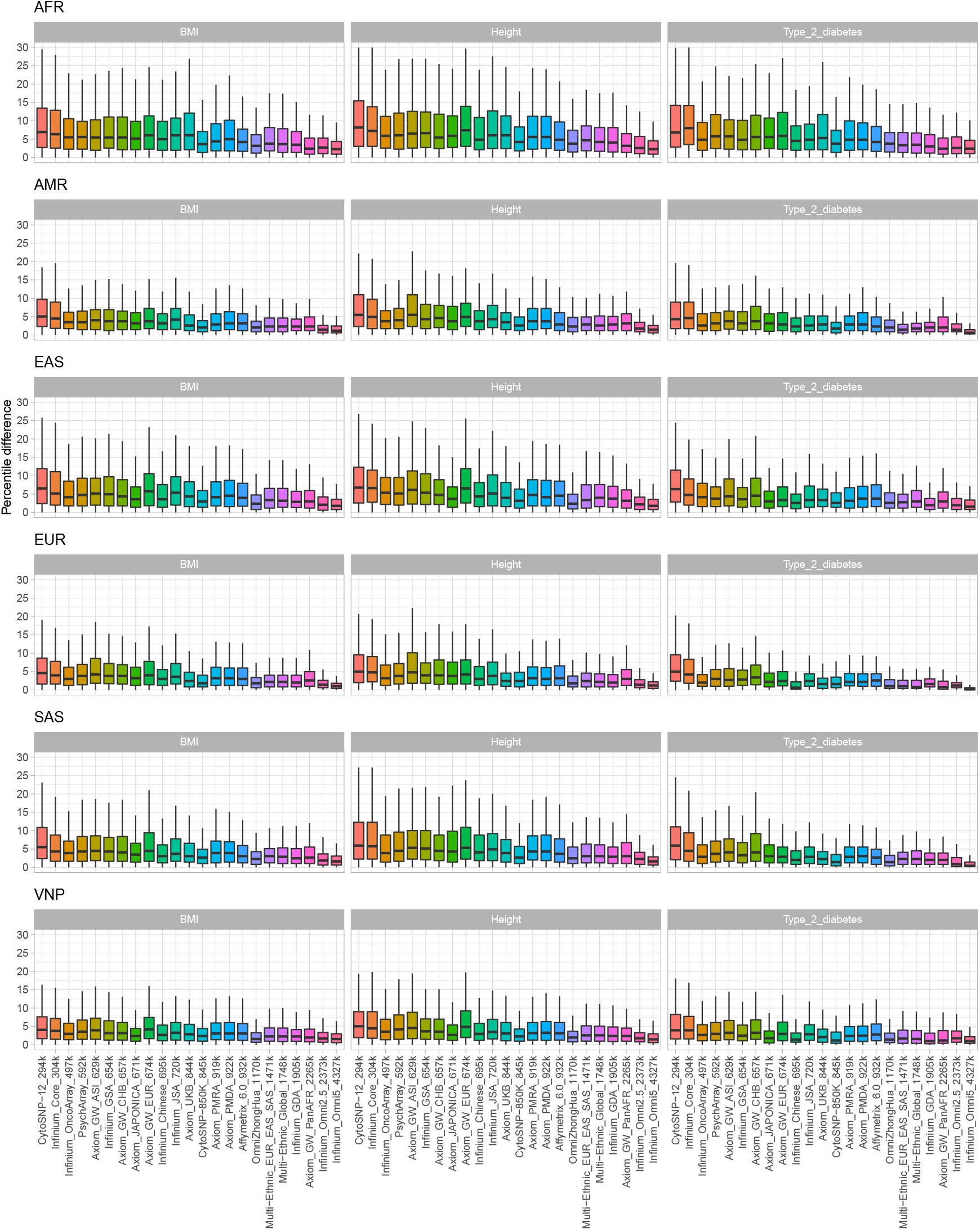
The absolute difference of percentile ranking between PGSs estimated from imputed genotyping data of 23 SNP arrays and PGSs estimated from WGS in six different populations. The figure shows results of three phenotypes including height, BMI, and type two diabetes with PRsice p-value setting of 5e-08.

In agreement with imputation performance, SNP arrays optimized specifically for targeted populations showed supperior PGS correlation in the targeted/closely related populations. For instance, Axiom Japonica Array NEO and Infinium OmniZhongHua v1.4 that was optimized for Japanese, and Chinese showed clear advantages in the populations of EAS, and VNP while Axiom UK Biobank Array yielded higher PGS correlation in the EUR population than the other size-comparable genotyping arrays. Taking height as a typical trait of interest, PGS correlations of the Japonica Array NEO were 0.9760, and 0.9847, while the Infinium OmniZhongHua v1.4 had 0.9879, and 0.9914 in EAS and VNP respectively. Interestingly, we observed that the Infinium CytoSNP-850K v1.2 was the array with superior PGS correlations in all populations, for all the three evaluated traits. For example, the PGS correlation for this array for height phenotype in AFR, AMR, EAS, EUR, SAS and VNP were 0.9679, 0.9876, 0.9789, 0.9908, 0.9844, 0.988, respectively.

Regarding the ADPR metric, the performance of arrays was in an agreement with the trend from comparing PGS correlation. ADPR in different PRSice-2 p-values settings are shown in Figure **??**, Figure S.2-12 and reported in detail in Table S.10-21. Most of the arrays yielded mean ADPR less than 10 in all three traits. An exception was the AFR populations with low density arrays. The highest density array, i.e. Infinium Omni5 v1.2, had the highest performance with ADPR less than 4. Notably, ADPRs varied by populations. Under-represented populations like AFR, and EAS tended to exhibit higher ADPRs than the others. Taking the p-value cutoff at 5e-8 for the height phenotype as an example (Figure **??**), Infinium Omni5 v1.2 obtained ADPRs of 3.8600, 2.4774, 2.8884, 1.9758, 2.8391, and 2,3699 in AFR, AMR, EAS, EUR, SAS and, VNP respectively. A consistent trend was also observed in other traits, with the lowest performance in AFR and the highest performance in EUR with ADPRs of 3.5974 and 1.8489 in BMI, and of 3.7206 and 1.6592 in type 2 diabetes. Similar to the other experiments, population specific arrays and the Infinium CytoSNP-850K v1.2 also illustrated their advantages when comparing the ADPR metric. The Axiom UK Biobank Array obtained good performance for the EUR population with ADPR of 3.0584, 3.1714, and 2.2734 in height, BMI, and type 2 diabetes respectively. This trend was also observed in the cases of Axiom Japonica Array NEO, and Infinium OmniZhongHua v1.4 applied for the EAS and VNP populations. Regarding the Infinium CytoSNP-850K v1.2 array, good performances in all traits and populations were observed. Specifically, ADPRs for height were 5.7141, 3.4914, 4.3753, 3.2501, 3.7638, 3.0267; for BMI at 4.9872, 2.5463, 4.1560, 2.6272, 3.5409, 3.1523; and for type 2 diabetes at 5.2000, 2.5762, 3.7687, 2.6066, 2.4707, 2.3812 in AFR, AMR, EAS, EUR, SAS and, VNP, respectively, all at the same p-value cutoff.

### 3.3 Performance evaluation using real genotyping data

We further utilize the availability of real genotyping data in the 1KVG dataset (95 out of the 1008 samples were also genotyped by Affymetrix PMRA array) to investigate how our simulated array data performed relative to the real array data. In brief, we simulated genotyping data of 95 samples by extracting variants from WGS data that matched with PMRA manifest, excluding phasing information. We then applied the same evaluation pipeline to compare the performance using simulated genotyping data against the results from the real genotyping data. In details, both simulated and real PMRA genotyping data were phased with SHAPEIT v4.1.3 (Delaneau et al., 2019), and imputed with Minimac4 v1.0.2 (Das et al., 2016). Reference data for both phasing and imputation were the remaining 913 WGS samples. Finally, imputation performance of both simulated and real PMRA arrays were estimated as described in the “Imputation performance evaluation” section. As expected, the imputation accuracies of simulated and real PMRA were highly concordant as shown in Figure 5 and Table 4. In details, mean and standard deviation of imputation accuracies of simulated PMRA are 0.7664±0.0309, 0.8522±0.0269, 0.9165±0.0179, and 0.9453±0.014; real PMRA are 0.7721±0.032, 0.8592±0.0278, 0.9217±0.0181, and 0.9497±0.0138 in four MAF bins of (0-0.01], (0.01-0.05], (0.01-0.5], and (0.05-0.5], respectively. These results indicated the robustness of our simulation approach for estimating imputation performances of genotyping arrays in reality.

**Table 4.** Mean and the standard deviation of imputation accuracies of simulated PMRA and real genotyped PMRA of 95 VNP samples at various MAF bins measured in 22 autosomes.

| MAF range | Simulated PMRA | Real PMRA |
| --- | --- | --- |
| (0-0.01] | $0.7664 \pm 0.0309$ | $0.7721 \pm 0.032$ |
| (0.01-0.05] | $0.8522 \pm 0.0269$ | $0.8592 \pm 0.0278$ |
| (0.01-0.5] | $0.9165 \pm 0.0179$ | $0.9217 \pm 0.0181$ |
| (0.05-0.5] | $0.9453 \pm 0.014$ | $0.9497 \pm 0.0138$ |

**Figure 5.**
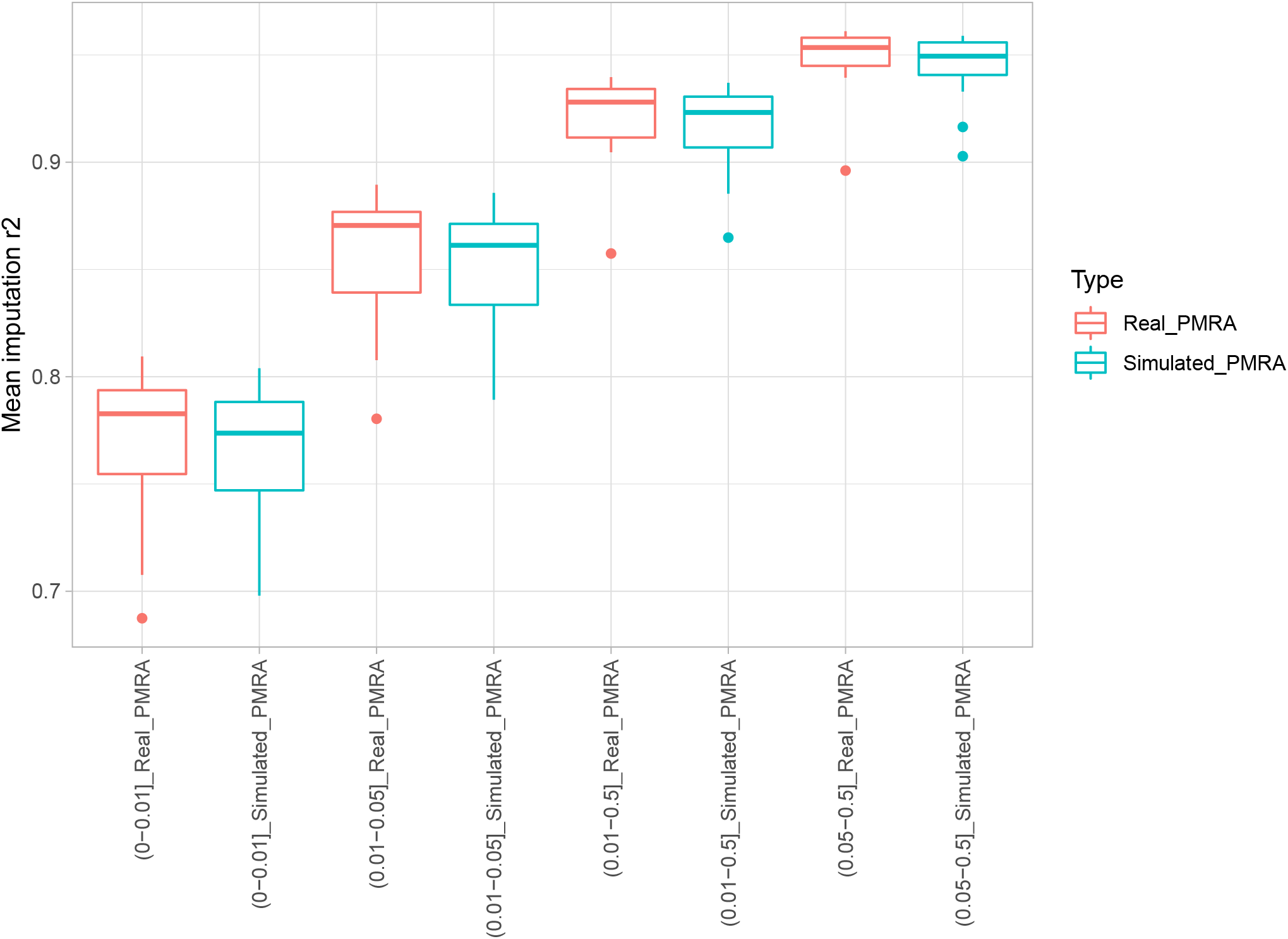
Mean imputation accuracy comparisons of simulated PMRA and real genotyped PMRA of 95 VNP samples at various MAF bins measured in 22 autosomes.

## 4 DISCUSSIONS AND CONCLUSIONS

Even in a booming time of next-generation sequencing technologies, current big genotyping projects are still using SNP arrays as the work-horse for generating valuable data, especially for bio-bank scale projects (Bycroft et al., 2018; Moon et al., 2019; Sakurai-Yageta et al., 2020). Moreover, genotyping by SNP arrays produce the exact information type required for PGS analysis, which is based on summarizing effect sizes from individual SNPs. A promising application of genomic research that is gaining increasing interest recently across the health-care system, and direct-to-consumer genomic services based on polygenic scoring like 23andMe (Lewis and Vassos, 2020; Folkersen et al., 2020). SNP arrays are clearly more economical in data generation and analysis, an important factor in designing project with a large sample size and/or with a limited budget. Given that there are many available human genotyping arrays optimized for various purposes, a comprehensive guideline for choosing the most suitable SNP arrays in multiple ancestry groups is still lacking. To address this gap, we have developed a systematic approach and tool to assess a large range of SNP arrays across multiple datasets. We performed imputation and PGS performance assessments for 23 human available genotyping arrays in six ancestry groups using both public and in-house datasets by various metrics. By comparing relative performance of SNP arrays relative to that of WGS with 4 metrics including imputation accuracy, imputation coverage, PGS correlation, and ADPR, we found expected trends and our results suggested suitable arrays that can maximize PGS performance for specific populations, especially for under-represented populations.

Overall, we found that all 23 assessed arrays had high performances in both imputation, and especially in PGS. These commercial arrays differ markedly in designs, i.e. the number of markers on the arrays and targeted ancestry groups that would cause performance deviations. An important finding in our analysis was that in order to obtain high imputation performances, the choice of array is not necessarily about getting higher density, but small to moderately-sized arrays (approximately 650k-850k tag SNPs), accompanied by well optimization for the targeted population could also produce high PGS performances. For example, the Japonica Array NEO, and the UK Biobank Array showed the highest performance when comparing with other arrays with the same sizes for EAS, and EUR populations respectively. This indicates that using customized, small-size SNP arrays at population-specific level can be a cost-effective genotyping solution without loosing PGS performance (Tam et al., 2019; Nguyen et al., 2022). We also observed that there are no specific arrays with moderate sizes that had superior imputation performances in AFR, and SAS, suggesting the need for genotyping arrays optimized for these populations. PGS performances were concordant to imputation performances in general. However, CytoSNP-850K v1.2 was an interesting array that showed superior PGS performances in all populations. This superior performance may be explained by the enrichment of cytogenomic regions in the design of the Infinium CytoSNP-850K v1.2 array (Illumina, ????). In agreement with previous studies (Martin et al., 2019; Sirugo et al., 2019), our analyses also show that underrepresented populations such as AFR, and SAS exhibited lower PGS performances (and ADPRs tended to be higher in AFR, and SAS) than other well-studied populations regardless the sample sizes of these populations are not significant different.

Notably, PGS performances of array constructed from imputed genotypes were very high in comparison with the original WGS PGS. The majority PGS correlations ranged from 0.90 to 0.99. In cases of optimal arrays for targeted populations in used (UK Biobank Array is used for EUR, Japonica Array NEO is used for EAS), the PGS correlation to WGS was higher than 0.97. In addition, PGS ranking differences between WGS and imputed array genotypes were not high with the majority of differences were under 5 percentile when optimal arrays were used. The possible reason for this observation was that current GWAS summary statistics were mostly generated by imputed array genotypes (Xue et al., 2018; Yengo et al., 2018) that are limited to detect rare associated markers. This indicates that using WGS for PGS analysis does not provide significant improvement interm of disease risk stratification at this time although this trend can change in the future when GWAS summary statistics at higher resolution become widely available (Wainschtein et al., 2022).

Finally, to make this analysis capability available to a broad audience, we developed a web application that provides interactive analyses SNP array contents and performances. As researchers may be interested in specific variants or regions, the application is aimed to support researchers to analyze SNP array contents and imputation performance based on population and genomic regions of interest. We hope that application will facilitate researchers in designing their genetic studies.

## CONFLICT OF INTEREST STATEMENT

There is NO Competing Interest.

## AUTHOR CONTRIBUTIONS

DTN initiated the study, designed experiments, analyzed data, interpreted results, developed the web tool, and drafted the manuscript. TT, MT, and NTD contributed to the 1KVG data generation and preprocessing. KT, DP, QN, and NSV contributed to the discussion, design and interpretation. QN and NSV critically revised the manuscript, coordinated the project, and supervised the study. All authors have read and approved the final manuscript.

## FUNDING

This work is funded by Vingroup Big Data Institute internal funding, and partly supported by the Vingroup Innovation Foundation under grant VINIF.DA.2020.02.

## ACKNOWLEDGMENTS

We especially thank Nguyen T. Nguyen for his kindly help in downloading the 1KGP-NYGC datasets, Mr Hoang H. Ho for the help with deploying the web tool. We also thank the Vingroup Big Data Institute for providing computational resources.

## DATA AVAILABILITY STATEMENT

The 1KGP-NYGC datasets are freely available at IGSR data portal (https://www.internationalgenome.org). The 1KVG WGS and PMDA array datasets are available under agreement at MASH data portal (https://genome.vinbigdata.org/). Data and source codes to generate figures of this study are available at: https://github.com/datngu/SNP_array_comparison. SNP array analyzing tool is available online at: https://genome.vinbigdata.org/tools/saa/. SNP-wise imputation performance estimation based on 1KGP-NYGC are freely available at: https://zenodo.org/record/6548098. SNP-wise imputation performance estimation based on 1KVG are available and can be supplied under ethical policy agreement.

## Notes

https://www.internationalgenome.org/

https://genome.vinbigdata.org/tools/saa/

https://zenodo.org/record/6548098

https://github.com/datngu/SNP_array_comparison

### Competing Interest Statement

The authors have declared no competing interest.

